# Assessment of the effects of aerobic fitness on cerebrovascular function in young adults using multiple inversion time arterial spin labelling MRI

**DOI:** 10.1101/539072

**Authors:** Catherine Foster, Jessica J Steventon, Daniel Helme, Valentina Tomassini, Richard G. Wise

## Abstract

The cross-sectional study investigated the effects of aerobic fitness on cerebrovascular function in the healthy brain. We quantified grey matter (GM) cerebral blood flow (CBF) and cerebrovascular reactivity (CVR), in a sample of young adults within a normal fitness range. Based on existing TCD and fMRI evidence, we predicted a positive relationship between fitness and resting GM CBF, and CVR. Exploratory hypotheses that higher 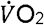 peak would be associated with higher GM volume and cognitive performance were also investigated.

20 adults underwent a 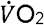 peak test and a battery of cognitive tests. All subjects underwent an MRI scan where multiple inversion time (MTI) pulsed arterial spin labelling (PASL) was used to quantify resting CBF and CVR to 5% CO_2_.

ROI analysis showed a non-significant negative correlation between whole-brain GM CBF and 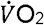 peak; r=-0.4, p=0.08, corrected p (p’) =0.16 and a significant positive correlation between 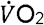 peak and voxelwise whole-brain GM CVR; r=0.62, p=0.003, p’ =0.006. Voxelwise analysis revealed a significant inverse association between 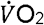 peak and resting CBF in the left and right thalamus, brainstem, right lateral occipital cortex, left intracalcarine cortex and cerebellum. The results of this study suggest that aerobic fitness is associated with lower CBF and greater CVR in young adults.

## Introduction

Neurodegenerative diseases are a global health burden and despite extensive research there are no cures or effective long-term management solutions. Aerobic fitness has emerged as a modifiable lifestyle factor which reduces the risk of all-cause mortality (Kodama et al., 2009; Lee, Artero, Sui, & Blair, 2010), cardiovascular events (Haskell, Lee, Pate, Powell, & Blair, 2007; Lee et al., 2010; Roque, Hernanz, Salaices, & Briones, 2013) and protects the brain against age and disease-related decline (Hayes, Alosco, & Forman, 2014; Hillman, Erickson, & Kramer, n.d.; Nishijima, Torres-Aleman, & Soya, 2016). In addition, brain structure and function are closely linked; to maintain healthy brain tissue, adequate energy must be supplied. Many studies have investigated the link between aerobic fitness and brain structure (Voss et al., 2015) but the relationship between aerobic fitness and cerebrovascular function has not been extensively studied in humans. The present study focuses on how aerobic fitness impacts the cerebrovascular processes which maintain delivery of brain nutrients.

Neuroimaging studies have provided evidence of a neuroprotective effect of aerobic fitness in older adults, linking higher aerobic fitness with greater brain volume (Erickson et al., 2011) and cerebral blood flow (CBF) (Thomas et al., 2013). Aerobic fitness has also been associated with greater cerebrovascular reactivity (CVR), which reflects the ability of the cerebral vessels to dilate, or the vascular reserve, assessed using transcranial Doppler (TCD) ultrasound (Bailey et al., 2013) in both older and young adults. However, Barnes et al. (2013), using the same technique, did not find a relationship between CVR and 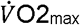 in younger adults despite observing the same trend as Bailey et al. (2013) in older adults. Further, the findings of Bailey et al. (2013) and Barnes et al. (2013) in older adults, do not agree with the MRI results of Thomas et al. (2013) who reported a lower BOLD-based CVR in masters athletes vs. sedentary young and age-matched controls.

Finally, Gauthier et al. (2015) found that in older adults, BOLD CVR was negatively correlated with 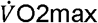 max in frontal regions but positively correlated in periventricular white matter and portions of the somatosensory cortex. These conflicting CVR data represent an issue of inconsistent findings regarding aerobic fitness and its effects on cerebrovascular function across the lifespan.

Recent findings from Hwang et al. (2018) using TCD, also suggest that young adults with higher levels of aerobic fitness have a greater cerebrovascular response to hypercapnia in the middle cerebral artery (MCA), during exercise, but found no differences at rest. The higher fit group also scored better on tests of fluid reasoning, but not on any other tasks included in the cognitive battery. However, as we are interested in how aerobic fitness may lead to general improvements in cerebrovascular health rather than vascular regulation during exercise, the current cross-sectional study focused on the relationship between aerobic fitness and cerebrovascular function in the resting brain of young, healthy adults. As the TCD literature suggests a positive relationship between fitness and CVR in young people, it is important to establish whether functional MRI reports the same effects, as TCD reflects the arteries rather than grey matter.

The objective of this study was to quantify variations in CBF and CVR, two widely used indices of cerebrovascular function, in a sample of young, healthy adults within a normal range of fitness levels (i.e. sedentary and active, but not athletes in structured training). Based on the extant TCD and fMRI evidence, we predicted a positive relationship between fitness and resting grey matter CBF. We also expected to see a positive correlation between CVR to CO_2_ and 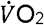 peak, a measure of aerobic fitness, in line with previous findings (Bailey et al., 2013; Hwang, Castelli, & Gonzalez-Lima, 2017). Tests chosen to measure a number of cognitive domains were included to examine the exploratory hypotheses that higher 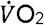 peak would be associated with better cognitive performance and higher grey matter volume. If CBF and CVR differences due to fitness were evident, then demonstrating additional cognitive benefits would provide evidence for functionally relevant effects of neurobiological differences. As the issue of inconsistent findings regarding CVR and 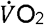 peak is potentially due to the different imaging methodologies employed, this study did not aim to resolve the issue completely. However, in using pulsed arterial spin labelling (PASL) MRI at multiple imaging delay times, we quantified global grey matter CBF and CVR in young adults. ASL provides quantitative assessment of global grey matter CBF unaffected by neuronal function, unlike BOLD fMRI and TCD respectively. Although ASL has SNR limitations, it is arguably the best technique to investigate CBF and CVR differences associated with 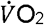 peak. The high labelling efficiency and lower sensitivity to off-resonance effects of PASL (Haller et al., 2016) make it well suited to acquisitions using CO_2_, as off-resonance effects can result in underestimation of perfusion where transit and arrival times differ between arteries.

## Methods

### Participants

20 healthy adults (11 females, mean age 25±4.6) were recruited from advertisements placed around Cardiff University. The study was approved by the Cardiff University School of Psychology Research Ethics Committee and performed in accordance with the guidelines stated in the Cardiff University Research Framework (version 4.0, 2010). All participants were non-smokers and educated to university level. Informed written consent was obtained from all subjects.

### Study Procedures

The study consisted of 3 lab visits. In Visit 1, eligibility screening for MRI, respiratory modulations (see figure S1) and intensive exercise was carried out. Contraindications to exercise were assessed using the Physical Activity Readiness Questionnaire (PARQ). Sociodemographic information was recorded and estimations of weekly activity level were established using the International Physical Activity Questionnaire (IPAQ) Short Form (Craig et al., 2003). Elevated CO_2_ inhalation can cause sensations of breathlessness, light headedness and anxiety in some individuals. For this reason, all volunteers took part in a gas modulation session in a MR scanner simulator. A stepwise protocol was employed to allow participants time to become accustomed to CO_2_ inhalation (see supplementary figure 1).

In Visit 2, volunteers completed 7 cognitive tests, administered by the same researcher for all participants, prior to the fitness test. The tests, covering a range of cognitive domains, were chosen as they are validated for use in patient and control samples (see table 1).

**Table 1.** List of cognitive tests used in this study and domains they are intended to assess.

| Test | Domain Measured |
| --- | --- |
| Speed and Capacity of Language Processing (SCOLP) (Baddeley, Emslie, Nimmo-Smith, 1992) | Information processing, speed and language comprehension |
| Forward Digit Span | Working memory |
| Letter Fluency (Categories) | Verbal fluency |
| Conners Continuous Performance Test (Conners et al., 2000) | Sustained attention and response inhibition |
| Trail Making Test (Part B) | Processing speed and executive function |
| Symbol Digit Modalities Test | Information processing speed |

**Table 2.** Group characteristics and fitness test outcomes including 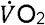 peak and secondary validation criteria. ETCO_2_ values represent values recorded during the ASL scan.

| Characteristics | Mean (sd) |
| --- | --- |
| Sex (11 female, 9 male) | - |
| Age (years) | 25 (4.6) |
| Weight (kg) | 69.1 (8.8) |
| Height (cm) | 173 (7) |
| BMI | 23 (2.1) |
| VO<sub>2</sub>peak (L/min) | 2.9 (0.6) |
| VO<sub>2</sub>peak (L/kg/min) | 41.2 (8) |
| Baseline Lactate mmol/L | 0.96 (0.3) |
| Post-Exercise Lactate | 9.5 (2) |
| Baseline HR | 79.2 (18.8) |
| Peak HR | 185.4 (9.8) |
| Baseline BP (pre- VO<sub>2</sub>peak) | 119/68 |
| Peak RER | 1.1 (0.04) |
| Maximum Work Rate (watts) | 205 (40) |
| Baseline Borg (legs) | 0.075 (0.2) |
| Peak Borg (legs) | 8.5 (1.7) |
| Baseline Borg (breathing) | 0.075 (0.3) |
| Peak Borg (breathing) | 7.55 (2) |
| Baseline ETCO<sub>2</sub> | 36.6 (3) |
| Hypercapnia ETCO<sub>2</sub> | 44.2 (3.8) |
| ETCO<sub>2</sub> increase during hypercapnia | 7.8 (1.4) |

### Fitness Test

The fitness test was also performed in Visit 2. The PAR-Q was conducted a second time to identify potential risk factors associated with exercise in the unlikely case of circumstance changes from Visit 1. The 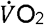 peak test protocol used has previously been described by (Collett et al., 2011; Cooney et al., 2013). The test began with a 2-minute unloaded warm up at 50 revolutions per minute (rpm) on a Lode cycle ergometer (Lode, Groningen, Netherlands). During the test, participants maintained a constant 50rpm and work rate was increased from 50 watts by 25 watts every 2 minutes. The test was terminated if any of the following criteria were reached; work rate fell below 45rpm for > 10 seconds, volitional exhaustion occurred, or maximal predicted heart rate exceeded 100%. At the end of each 2-minute step Borg ratings of perceived exertion, on the CR10 scale (Borg & Kaijser, 2006) were used to record perceived heaviness of legs and breathing.

Blood lactate concentration was sampled from the earlobe using Unistik 3 1.8mm lancets (Williams Medical, Caerphilly, Wales) and tested using the Lactate Plus system (Nova Biomedical, Waltam, MA USA) at baseline, 2-minute intervals and at exhaustion. Blood pressure (BP) was recorded at baseline, HR at baseline and 2-minute intervals during the test. Respiratory gas exchanges were continuously monitored using breath by breath analysis with the Cortex Metalyser 3b (Cortex Biophysik Metalyzer, Germany). 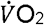 peak was taken as the highest value over a 30 second block average at the maximum recorded work rate. Heart rate (HR) and work rate were also recorded at baseline and at 2-minute intervals and electrocardiography (ECG) was monitored for the duration of the test. Following termination of the 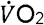 peak test participants were monitored for 10-minutes. In this 10-minute recovery period blood pressure, HR and blood lactate were sampled every 2 minutes before subjects left the lab.

### MRI Acquisition

In Visit 3, volunteers underwent the MR scan. Images were acquired on a 3T whole body MRI system (GE Excite HDx, Milwaukee, WI, USA) using an eight-channel receive-only head coil. Heart rate was recorded using a pulse oximeter and a respiratory bellows monitored breathing. A sampling line connected to the face mask of the breathing circuit(Felipe B Tancredi, Lajoie, & Hoge, 2014) was used to monitor the partial pressure of end-tidal CO_2_(P_ET_CO_2_) and O_2_ (P_ET_O_2_) via the Biopac_®_ system (Biopac, Worcestershire, UK). The MEDRAD system (MEDRAD, Pittsburgh, PA) was used to monitor O_2_ saturation throughout the experiment.

PASL data were acquired using a single subtraction PICORE QUIPSS II (Wong et al., 1998) with a dual-echo gradient-echo readout (Liu et al., 2002) and spiral k-space acquisition (Glover, 1999) the first echo being used for CBF quantification. Data were acquired at 8 inversion times; 400, 500, 600, 700, 1100, 1400, 1700 and 2000ms. QUIPSS II cut-off at 700ms meant that short and long inversion times were acquired in separate runs. 16 and 8 tag-control pairs were acquired for each of the short and long inversion time respectively. A variable TR was used for efficiency, 1200-1500ms for short TI data (400-700ms) and 1700-2600ms for long TIs (1100-2000ms). Other acquisition parameters were; TE = 2.7ms, voxel size = 3.5×3.5×7mm_3_, 3.2mm in-plane resolution, matrix size 64×64mm, FOV = 19.8cm, flip angle = 90°, slice delay 55ms, 15 slices, slice gap 1.5mm for maximum brain coverage. Label thickness was 200mm with 10mm gap between the end of the label and the most proximal imaging slice.

A calibration image without any labelling was acquired before the perfusion mages using the same acquisition parameters, except a long TR; this was used to obtain the equilibrium magnetisation of cerebrospinal fluid (M_0_, CSF), needed for the quantification of CBF. A minimal contrast image was also acquired to correct for coil inhomogeneities with TE = 11ms, TR = 2s. The PASL acquisition was performed twice, first during rest, then while volunteers were hypercapnic. The hypercapnic modulation was carried out using the prospective control method described by (Tancredi & Hoge, 2013). A gas mixing chamber constructed in-house had three feeding lines coming in for the delivery of medical air, 5% CO_2_, and medical oxygen, the latter incorporated as a safety backup but not used during experimentation, the circuit is described in detail by (Tancredi, Lajoie, & Hoge, 2014). P_ET_CO_2_ elevation of 7mmHg was targeted using 5% CO_2_ (balance air) and the scan commenced once participants reached the target CO_2_ level. ASL data acquisition lasted ∼6 minutes during rest, followed by a delay to allow the onset of hypercapnia. Once a P_ET_CO_2_ elevation of approximately 7mmHg was achieved, the ∼6 minute ASL acquisition was repeated.

A T1-weighted 3D structural fast spoiled gradient echo (FSPGR) scan was also acquired for registration of functional data and voxel-based morphometry (VBM) analysis; TR/TE = 7.8/2.9ms, resolution = 1mm isotropic.

## Data Analysis

### 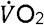 peak Calculation

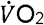 peak was calculated from the respiratory gas exchanges which were continuously monitored using breath-by-breath analysis and averaged over 10 second blocks. 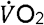 peak was defined as the 30-second averaged value at maximum recorded work rate. HR, respiratory exchange ratio (RER) and blood lactate were recorded every 2 minutes and at termination of the fitness test to validate 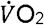 peak criteria being reached. The test result was considered valid if heart rate exceeded 90% of age predicted maximum, RER >1 and blood lactate concentration was greater than 8 mM at the end of the test (Horton et al., 2011; Milani, Lavie, Mehra, & Ventura, 2006; Poole, Wilkerson, & Jones, 2008).

### Cognitive Data Scoring

Cognitive data were scored by hand by the same researcher, the Conners Continuous Performance test is computer based and scored automatically. Demeaned scores were calculated for correlation analysis with 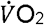 peak and MR data.

### Image Analysis

Physiological noise correction was carried out using a modified RETROICOR (Glover, Li, & Ress, 2000) to remove cardiac and respiratory noise components from the data. ASL data were motion corrected using MCFLIRT within FSL (FMRIB’s Software Library, www.fmrib.ox.ac.uk/fsl, (Jenkinson, Bannister, Brady, & Smith, 2002). The CBF timeseries for the normocapnia and hypercapnia scans were corrected for coil sensitivity inhomogeneities using the minimal contrast images (Wang, Qiu, & Constable, 2005). Average difference images were obtained for each inversion time from tag-control subtraction of the CBF time series (Lu, Donahue, & Van Zijl, 2006) and a perfusion map was created from all inversion times using a two-compartment CBF kinetic model implemented using the BASIL toolkit of FSL (Chappell et al., 2010). Within this framework, perfusion maps were converted to ml/100g/min using the CSF signal as a reference to estimate the fully relaxed magnetisation of water in blood (Chalela et al., 2000). The mean resting CBF for each subject was calculated by averaging the CBF time series over all time points and over all voxels within the masked grey matter image (see below). CVR was calculated by dividing the percentage change in CBF from rest during hypercapnia by the mmHg change in P_ET_CO_2_ increase during hypercapnia.

### Grey Matter Volume

T1-weighted structural data were analysed with FSL-VBM (Douaud et al., 2007; Good et al., 2001) http://fsl.fmrib.ox.ac.uk/fsl/fslwiki/FSLVBM). Structural images were brain extracted and segmented into grey matter, white matter and CSF before being registered to the MNI 152 standard space template using non-linear registration (Andersson et al., 2007). The resulting images were averaged and flipped along the x-axis to create a left-right symmetric, group-specific GM template. Second, all native GM images were non-linearly registered to this study-specific template and modulated to correct for local expansion (or contraction) due to the non-linear component of the spatial transformation. The modulated GM images were then smoothed with an isotropic Gaussian kernel with a sigma of 3mm. Finally, voxelwise GLM was applied using permutation-based non-parametric testing, correcting for multiple comparisons across space to investigate whether GM volume and 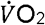 peak were correlated.

### ROI Analysis of Whole Grey Matter CBF and CVR

Individual subject grey matter masks were created using FMRIB’s Automated Segmentation Tool (FAST) and thresholded to include voxels with >50% grey matter probability. ROI analysis of the relationship between grey matter CBF at rest, CVR and 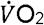 peak was carried out using correlation analysis and permutation testing (100,000 permutations per comparison) in Matlab (Mathworks Inc., MA, USA). Permutation testing was used as an alternative to Bonferroni correction for multiple comparisons which is too conservative when potentially related variables are tested. This approach was also applied to examine the associations between grey matter CBF and CVR, and cognitive test scores.

### Voxelwise Analysis of Grey Matter CBF and CVR

Follow up voxelwise analysis was conducted using FSL’s Randomise tool (http://www.fmrib.ox.ac.uk/fsl/randomise/index.html) (Winkler, Ridgway, Webster, Smith, & Nichols, 2014) to examine the voxelwise correlation between i) CBF and 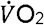 peak and ii) CVR and 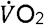 peak. The individual subject masks described in the ROI Analysis section were used to confine this analysis to global grey matter only. Randomise is a permutation testing method that uses threshold free cluster enhancement (TFCE) (Smith & Nichols, 2009) to correct for multiple comparisons across voxels. Significance was set at p < 0.05 (FDR corrected).

## Results

### Demographics and Physiological Information

All subjects completed the fitness test. Out of the 20 subjects 17 achieved an RER > 1.1 and 3 achieved RER > 1. 17 subjects exceeded the 90% HR maximum threshold and achieved a lactate peak >9, the remaining 3 exceeded 80% of HR maximum and achieved a lactate peak >6. As this was a 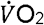 peak test all subjects satisfied test criteria by working to volitional exhaustion. Based on typical 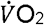 max criteria, 17 subjects fulfilled criteria for a maximal exercise test (Dupuy et al., 2015; Wilkinson, Leedale-Brown, & Winter, 2009). No adverse effects due to elevated CO_2_ inhalation during the MRI scans were reported in this sample.

### Correlations between CBF, CVR and 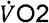 peak

#### ROI Analysis

There was a non-significant inverse association between 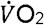 peak and resting CBF in grey matter; r = −0.4, p = 0.08 p’ =.16 (figure 1) and a significant positive correlation between 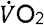 peak and CVR; r = 0.62, p = 0.003, p’ =0.006 (figure 2).

**Figure 1.**
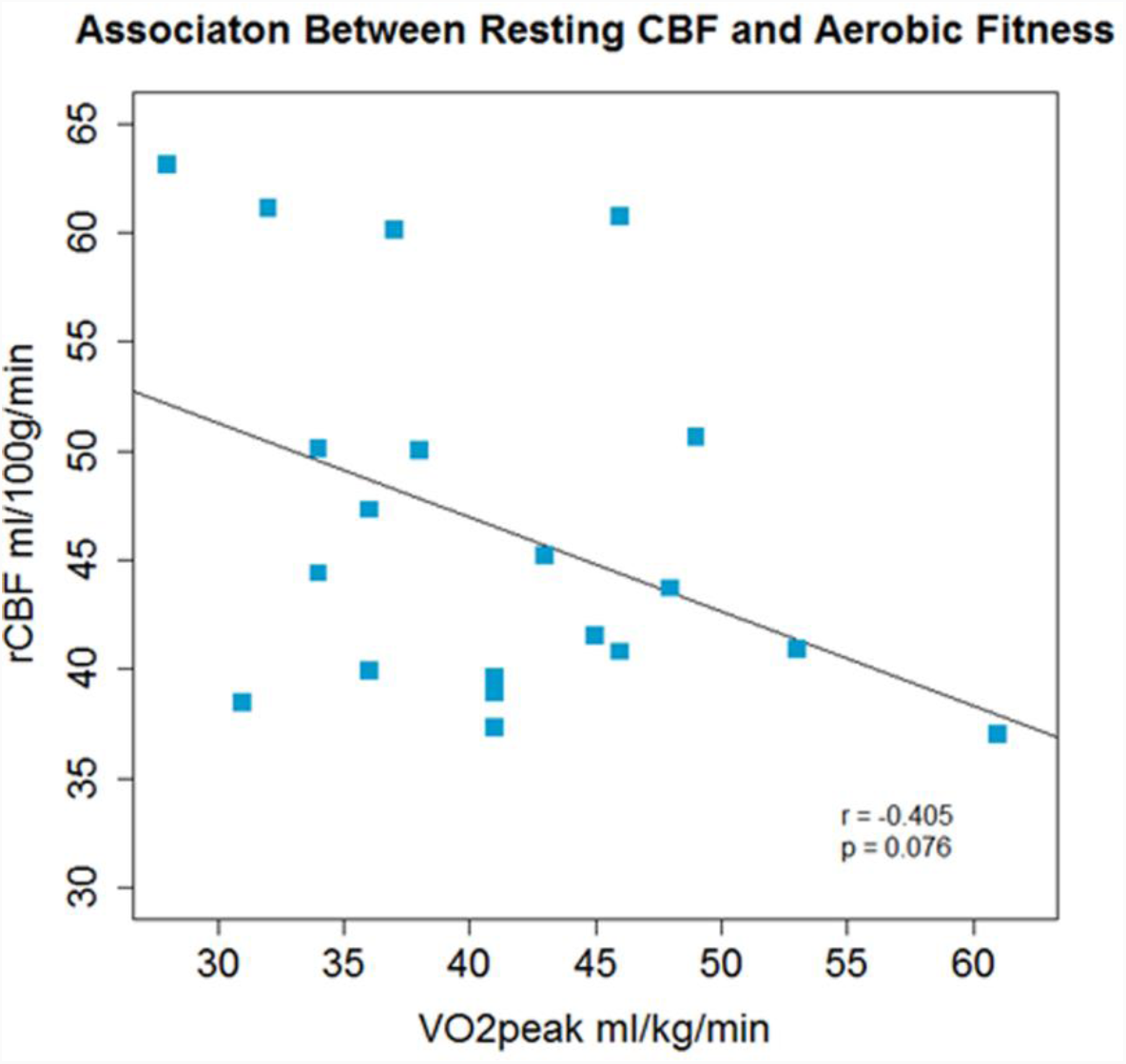
A non-significant inverse association between aerobic fitness and whole brain grey matter CBF was observed.

**Figure 2.**
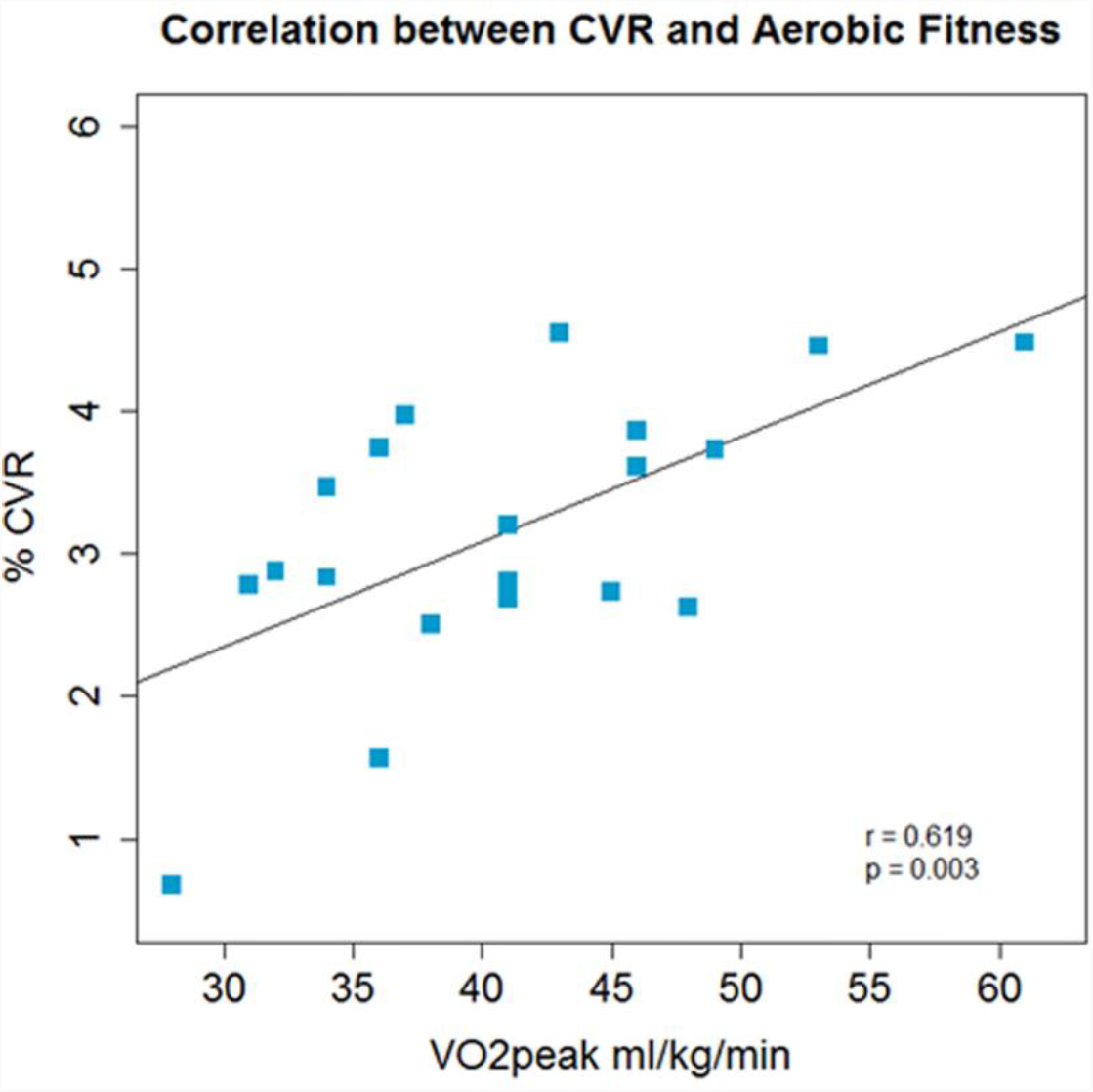
Across the group whole brain grey matter CVR and aerobic fitness were positively correlated.

Follow-up exploratory analysis showed that baseline P_ET_CO_2_ was not associated with 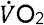 peak; r = −0.05, p = 0.8 (supplementary figure 2), nor was the P_ET_CO_2_ change during hypercapnia associated with 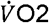 peak; r= 0.26.p = 0.27.

#### Voxelwise analysis

Voxelwise analysis revealed a significant inverse association between 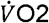 peak and resting CBF in the left and right thalamus, brainstem, right lateral occipital cortex, left intracalcarine cortex and cerebellum (figure 3). No regions were significantly positively correlated with 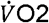 peak. Voxelwise CVR did not correlate with 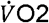 peak.

**Figure 3.**
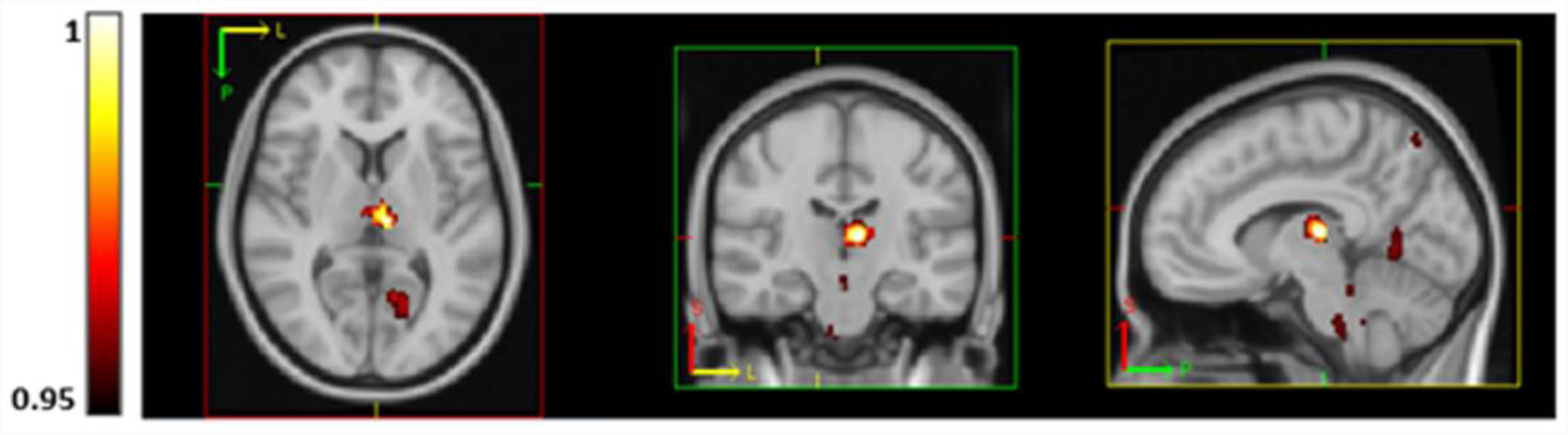
Regions of significantly lower CBF (using TFCE thresholding and FEW corrected) in subjects with higher aerobic fitness at rest in the thalamus, brainstem, precuneus, visual cortex (V1) and lingual gyrus. P-values are displayed as 1-p where a value of 1 is most significant.

#### Cognitive Performance, Grey Matter Volume and 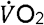 peak

Finally, correlation analysis did not reveal any significant associations between 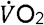 peak and cognitive performance, or cognitive performance and cerebrovascular measures. Mean scores and correlations are shown in supplementary tables 1 and 2. VBM analysis of grey matter volume did not show a trend or association with 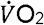 peak.

## Discussion

### Summary

In this study, we examined the association between aerobic fitness and CBF and CVR using multiple inversion time PASL. This is the first study to report associations between aerobic fitness, CBF-based CVR as opposed to BOLD-based CVR, and whole grey matter CBF in young adults using ASL. Across the group, higher 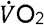 peak was associated with lower regional resting CBF and greater global grey matter CVR to CO_2_. 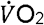 peak was not associated with cognitive task performance or grey matter volume in this cohort.

### 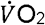 peak, CBF and CVR

The inverse correlation between CBF and 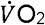 peak was not expected, as the opposite has been found in children (Chaddock-Heyman et al., 2016), young adults (Bailey et al., 2013) and older adults (Ainslie et al., 2008; Bailey et al., 2013; Brown et al., 2010; Thomas et al., 2013; Zimmerman et al., 2014). However, with the exception of Thomas et al., (2013), Zimmerman et al., (2014) and (Chaddock-Heyman et al., 2016), previous studies have used TCD ultrasound. TCD measures blood flow velocity, not flow, as TCD does not measure vessel diameter which would be required to calculate flow using this method. In contrast, ASL measures CBF directly through the labelling of inflowing water to tissue. Therefore, results may differ as they are measuring different, albeit related, vascular parameters. Recent efforts to characterise differences between MRI and TCD measures of CVR are valuable and will allow the methods to be used as complementary techniques given that TCD can be conducted under conditions not suited to MRI. For example, Burley et al. (2017) reported a positive correlation between VO_2_max and CVR using both BOLD fMRI and TCD suggesting that the methods are not necessarily contradictory (Bailey et al., 2013; Thomas et al., 2013) and could be used to investigate brain function within groups at rest and during exercise. The main limitations of TCD are that it lacks the spatial resolution and range of neurophysiological measures that can be recorded with quantitative MRI.

Most existing MRI studies investigated groups undergoing neurodevelopment (Chaddock-Heyman et al., 2016) and older adults (Thomas, Liu, Park, van Osch, & Lu, 2014; Zimmerman et al., 2014) where the mechanistic relationships with aerobic fitness may be different to adults not in stages of either development or decline which may explain why results are not consistent across studies with different age groups

In this study, CBF showed a non-significant inverse correlation with 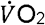 peak across the whole of grey matter. Statistical significance was localised to regions of the thalamus, brainstem, visual cortex, cerebellum and precuneus. Regional variation in contrast-to-noise may explain why these specific regions correlated with fitness in this sample, however, previous studies have also reported regional rather than global effects.

A study by Robertson et al. (2015) in stroke survivors, using single inversion time PCASL, similarly found an inverse correlation between precuneus CBF and aerobic fitness, but a positive correlation between thalamic and posterior cingulate CBF and aerobic fitness. Gauthier et al. (2015) found both positive and inverse correlations between CVR and aerobic fitness in older adults, and this, along with the Robertson et al. (2015) study adds to the evidence that aerobic fitness may have complex, regionally varying and age dependent effects on the brain. Although the direction of the correlation in stroke survivors in the thalamus and posterior cingulate cortex contradicts the current results, the current study and those of Robertson et al. (2015) and Gauthier et al. (2015) further suggest that aerobic fitness may exert localised effects on CBF (Chaddock-Heyman et al., 2016; Thomas et al., 2013). Given that participants in each of these cited studies have differed in age and health status, the fact that consistent findings have not been reported is unsurprising. The variance in the effects of aerobic fitness on brain function with age needs further investigation through interventions in young and older groups as there may be a cross-over effect whereby higher CBF in youth and older adulthood has different functional importance. The duration of a given fitness level may also be important in interpreting results; for example, those who have maintained a high fitness level throughout their life may have a different cerebrovascular profile from those who more recently achieved a similarly high 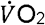 score. The current study consists of a normal sample of young adults who varied in fitness level, but not to the extent that a cross-sectional sample comprised of athletes and sedentary individuals would. However, our sample had a mean 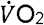 peak of 41 ml/kg/min, standard deviation 8, similar to Hwang et al., (2017) who reported an average 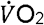 max of 37.8, standard deviation 9.6 in a sample of 87 participants and also found a positive correlation between aerobic fitness and CVR in the MCA. The range of fitness levels within the study sample is likely to dictate the degree of observed fitness-related brain effects, especially in healthy groups. In fact, as well as differences in study design, the spread of fitness levels within studies may also explain the differences in the results of Barnes et al. (2013) and Bailey et al. (2013). Bailey et al. (2013) compared CVR between two groups, sedentary vs. trained, whereas Barnes et al. (2013) used a single group with a range of fitness levels who did not participate in a structured training regime.

There is a neurobiological interpretation which potentially explains the negative correlation between CBF and 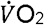 peak in this study. Regular exercise training which increases 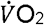 max, can result in increased tissue capacity to extract oxygen from blood in muscle (Kalliokoski et al., 2001; Ookawara et al., 2002), potentially as a result of greater blood vessel density which facilitates oxygen diffusion throughout the tissue. A higher cerebral oxygen extraction fraction (OEF) in more aerobically fit subjects could explain the negative correlation observed, as less CBF would be required to maintain the cerebral rate of oxygen metabolism. In diseases where hypoperfusion is present, OEF is higher, due to the inverse correlation between CBF and OEF (Germuska et al., 2019), and cerebral autoregulation works to maintain oxygen supply to the brain when CBF is suboptimal.(Derdeyn et al., 2002; Román, Erkinjuntti, Wallin, Pantoni, & Chui, 2002). However, in the healthy brain, assuming SaO_2_ is close to 1, greater fitness-related OEF may be an indirect result of higher oxygen availability in the brain allowing it to extract the required energy more efficiently and thus a lower rate of CBF is sufficient. In later adulthood, this efficiency may serve to sustain cerebrovascular function meaning that optimal blood flow is maintained in trained subjects, as the typical age-related CBF decrease is offset. This could explain the seemingly opposing effects of higher aerobic fitness on CBF seen in this study in young adults and existing work in older adults (Chapman et al., 2013; Thomas et al., 2013). Further evidence for differential effects of aerobic fitness with age comes from a recent structural MRI study by Williams et al. (2017) which reported a positive correlation between cortical thickness and 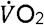 peak in older adults but an inverse correlation between these measures in young adults. In terms of function and metabolism, future studies examining the relationship between aerobic fitness, CBF and OEF would clarify whether the evidence supports this explanation. CBF quantification in older adults must also be interpreted with caution where partial volume effects have not been controlled for. Cortical atrophy and delayed bolus arrival times due to ageing combined with ASL voxel sizes of ∼7mm mean that perfusion may be underestimated where partial volume correction is not carried out.

Data from experimental models supports the current results. Young animals subjected to wheel running for several weeks had increased angiogenesis, and therefore capillary density, compared with sedentary animals (Pereira et al., 2007; Swain et al., 2003). A similar process may occur in humans where capillary density is higher in those with greater aerobic fitness as a result of long-term exercise participation, and in turn, such individuals may have greater oxygenated blood availability without necessarily increasing CBF. Lower capillary density would restrict the diffusion of oxygen into tissue as the oxygen gradient would be lower (Leithner & Royl, 2013), therefore, higher CBF rates under certain conditions may reflect vascular inefficiency where greater flow rates are necessary for optimal brain oxygenation. However, the short duration of interventions and lifespan of rodents means that the increased capillary density seen in these studies may not be comparable to work investigating effects of long term aerobic fitness, gained through years of physical training in humans.

Upon neuronal activation, there is an increase in demand for energy; in the healthy brain this is met with an increase in CBF. The lower resting CBF observed in higher fit individuals, although initially counterintuitive, is not problematic if there is an adequate vascular reserve to meet changes in energy demand. The data support this argument as a greater grey matter CVR to CO_2_ was observed in subjects with a higher 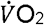 peak. If aerobic fitness maintains CBF and oxygenation, both of which are affected by ageing (Aanerud et al., 2012), then an increase in CBF, and potentially CVR, in the trained group, would be expected when comparing trained versus sedentary older adults. However, if aerobic fitness also enhances oxygen extraction efficiency, in young people this so-called efficiency may be observed as a reduced resting CBF if aerobic fitness does indeed increase the number of blood vessels and therefore CBV as outlined above. Greater vessel density would increase the capacity for oxygenation and oxygen diffusion gradient across the tissue. In addition, CVR and 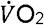 peak may then be positively correlated due to a higher vascular reserve, leading to a greater CBF response to CO_2_ as CBF increases well above what is required to respond to a stimulus (Raichle, 1986). Future work which quantifies OEF, CBF and CVR in similar populations could provide the additional necessary information to understand cerebral vascular and metabolic function at different levels of aerobic fitness.

The ROI analysis showed a positive correlation between CVR and 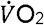 peak in global grey matter, however no specific regions emerged in a follow-up voxelwise analysis. CVR declines with age (De Vis et al., 2015; Peng et al., 2018) but in the literature, there are contradictory reports on the association between CVR and aerobic fitness (Bailey et al., 2013; Barnes, Taylor, Kluck, Johnson, & Joyner, 2013; Gauthier et al., 2015; Thomas et al., 2013), as discussed in the introduction. The positive association between 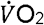 peak and CVR reported here is in line with the TCD findings by Bailey et al. (2013), whereby trained young adults had greater CVR than a sedentary comparison group. However, few studies have investigated the relationship between CVR and aerobic fitness in healthy young adults, therefore more work is needed to adequately understand this relationship. Overall results across studies using MRI and TCD to study CBF, flow velocity and CVR suggest that exercise training and aerobic fitness has complex and varying effects on brain regions with potential mechanistic differences across the lifespan. This complexity presents an important area of future research in different age groups, and a need to comprehensively map cerebrovascular function across the brain.

Finally, it should be noted that De Vis et al. (2015) showed that resting P_ET_CO_2_ accounted for age-related CBF differences almost entirely. In this study, there was a non-significant positive trend (r = 0.4, p = 0.06) between baseline P_ET_CO_2_ and CBF, but no strong associations between 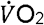 peak and baseline P_ET_CO_2_(r=0.05, p=0.8) or the ΔP_ET_CO_2_ during hypercapnia (r=0.26, p = 0.27, see supplementary figures S2-s4). This suggests a cerebrovascular effect of fitness on CVR rather than an effect driven by P_ET_CO_2_ differences.

### Cognitive Performance and Grey Matter Volume

Cognitive performance scores were similar across the group and there were no strong associations with 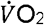 peak nor with CBF or CVR (see supplementary tables 1 and 2). It is possible that cognitive differences in young adults may only be visible at more extreme ends of the fitness spectrum than sampled in this study. It is also possible that more sensitive cognitive tests are required to detect differences among high functioning groups separated only by fitness level.

The cognitive reserve hypothesis (Chodzko-Zajko & Moore, 1994) states that higher fitness in older age offsets age-related decline in cerebral circulation, enhancing oxygen delivery to support neural demand. More recently, (Etnier, Nowell, Landers, & Sibley, 2006) conducted a meta-regression on studies examining the effects of fitness on cognitive function. In summary, the authors did not find strong support for a beneficial effect of fitness in any age group once moderator variables were considered, and in fact found a negative association between fitness and cognition in studies using a pre-post design. Only a very small number of studies report a ebneficial effect of fitness or physical activity on select cognitive domains in young people (Baym et al., 2011; Themanon, Pontifex & Hillman, 2009; Stroth, Hille, Spitzer, & Reinhardt, 2016 and other studies report significant effects in older but not young adults (Hayes, Forman, & Verfaellie, 2014). In summary, more work is needed in young adults to determine neurophysiological effects of aerobic fitness and their functional relevance in terms of health and cognition.

Aerobic fitness is believed to reduce age-related atrophy, mainly in the hippocampus (Erickson et al., 2011; Firth et al., 2018). In this study the focus was on cerebrovascular function, not brain structure, however, if structural differences were present, this would affect CBF and CVR. The VBM analysis showed that grey matter global volume differences due to fitness were not present. As this group was young and healthy, this finding is not wholly surprising, and suggests that volumetric differences only become apparent in later life, adding to our knowledge of the lifelong effects of aerobic fitness.

## Limitations

Haemoglobin (Hb) levels were not measured. Hb is a protein molecule found in red blood cells which is responsible for transport of O_2_ to tissue. The concentration of Hb in blood affects exercise performance; reduced Hb means that blood can carry less oxygen (Calbet, 2006) and therefore muscle function is impaired. In addition, Hb affects perfusion estimates as there is an inverse relationship between [Hb] and the longitudinal relaxation time (T_1_) of blood. Brain Hb cannot be measured directly *in-vivo*, however capillary samples may provide an indication of Hb differences between subjects or groups which could help to explain biological mechanisms driving CBF differences.

Second, investigations were restricted to grey matter. Due to the limited SNR of ASL, reliable quantification of CBF and CVR in white matter is difficult as CBF is much lower than in grey matter. However, adequate blood supply and energy metabolism is necessary for all round brain health, and ageing is associated with damage to white matter microstructure and reductions in myelination (Gunning-Dixon, Brickman, Cheng, & Alexopoulos, 2009). Therefore, improvement of methods to study the WM vasculature are also needed to understand the global effects of fitness.

The voxelwise and ROI analyses did not show the same statistical significances. While the voxelwise analysis of CBF revealed regions of significantly lower CBF with fitness, the ROI analysis showed a non-significant trend in the same direction for global grey matter. The ROI analysis of CVR however, showed a moderate (0.6) significant correlation with fitness while the voxelwise analysis did not reach statistical significance. ASL is an intrinsically low SNR technique compared to BOLD fMRI. Although ASL offers significant benefits in terms of quantification of physiological parameters, the SNR may have prevented detection of greater voxelwise associations with 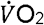 peak.

Lastly, there are known hormonal effects on exercise test performance (Jonge, 2003) and CBF (Brackley, Ramsay, Pipkin, & Rubin, 1999; Duckles & Krause, 2007). In the present study, the resting CBF difference was not significantly different between males and females; t(18)=-0.602, p = 0.094, but controlling for menstrual cycle phase, which was not done here, could reduce variability within cohorts. This is a preferable option to limiting studies to males only as hormonal differences may play a role in the acute response to exercise and possibly in mediating the effects of fitness on brain health.

### Summary and Future Directions

The results of this study suggest that aerobic fitness is associated with lower CBF and greater CVR in young, healthy adults, however the modest effects observed need replication in larger samples. We currently know very little about the functional relevance of CBF or CVR differences in young adults, and how the observed neurophysiological effects of physical training differ from those observed in older adults. With recent advances in quantitative MRI techniques, non-invasive mapping of multiple indices of cerebrovascular function e.g. CBF, CVR, CBV OEF and the cerebral metabolic rate of oxygen consumption is possible within a single scan session (Bulte et al., 2012; Gauthier, Desjardins-Crépeau, Madjar, Bherer, & Hoge, 2012; Gauthier & Hoge, 2013; Wise, Harris, Stone, & Murphy, 2013). Notably, regional CBV quantification would bring us a step closer to comparisons with experimental data on angiogenesis and capillary density following exercise as CBV and capillary density are presumed to be linked.

Application of these techniques to study brain function in both young and older trained and sedentary adults will provide information necessary to move forward in developing exercise training protocols to increase the adoption of fitness training as a preventative health tool. Also, although complete standardisation of data acquisition and analysis pipelines among researchers in this area is not realistic or warranted given the variety of questions to be addressed, a degree of standardisation would be helpful in resolving the differences found in the literature between MRI studies of CVR (Gauthier et al., 2015; Thomas et al., 2013). Further, making data freely available, so that researchers can pool data from similar studies using ASL or BOLD to investigate the cerebral effects of fitness, would build a clearer picture of effects across the age and fitness level spectrums. Importantly it would also increase statistical power which is sometimes lacking in MRI studies due to the high cost of the technique.

Research over the next decade should also work to establish whether regular exercise regardless of intensity level delivers brain benefits, or whether there is an aerobic fitness threshold, above which benefits such as maintained CBF with ageing, are observed. Answering this question will guide optimal exercise dose recommendations and interventional studies.

## Conflicts of interest

The authors do not declare any conflicts of interest.

## Authors’ contributions

Study design (CF, JJS, RGW), data collection (CF, JS, DH), data analysis (CF, RW), manuscript preparation (CF, VT, RW).

## Supplementary Data

**Figure S1.**
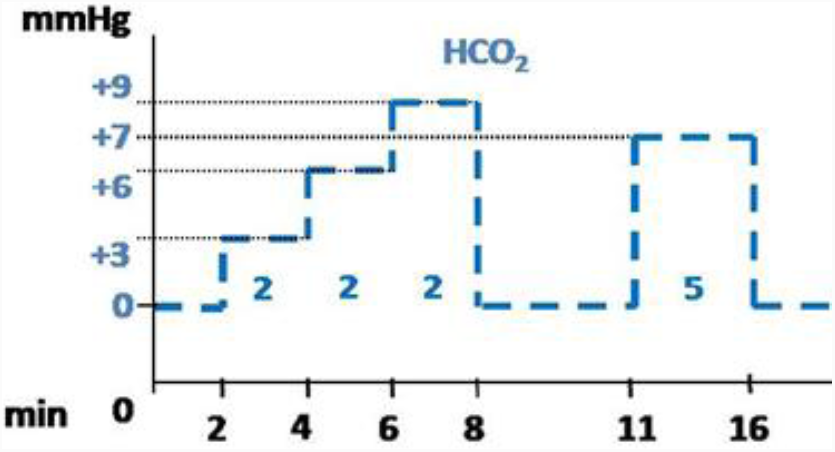
Protocol for the pre-MRI respiratory modulation test session. Heart rate and oxygen saturation were monitored and recorded throughout the session. Figure adapted from Merola, (2016). HCO2 = End-tidal CO2 during hypercapnia.

**Figure S2.**
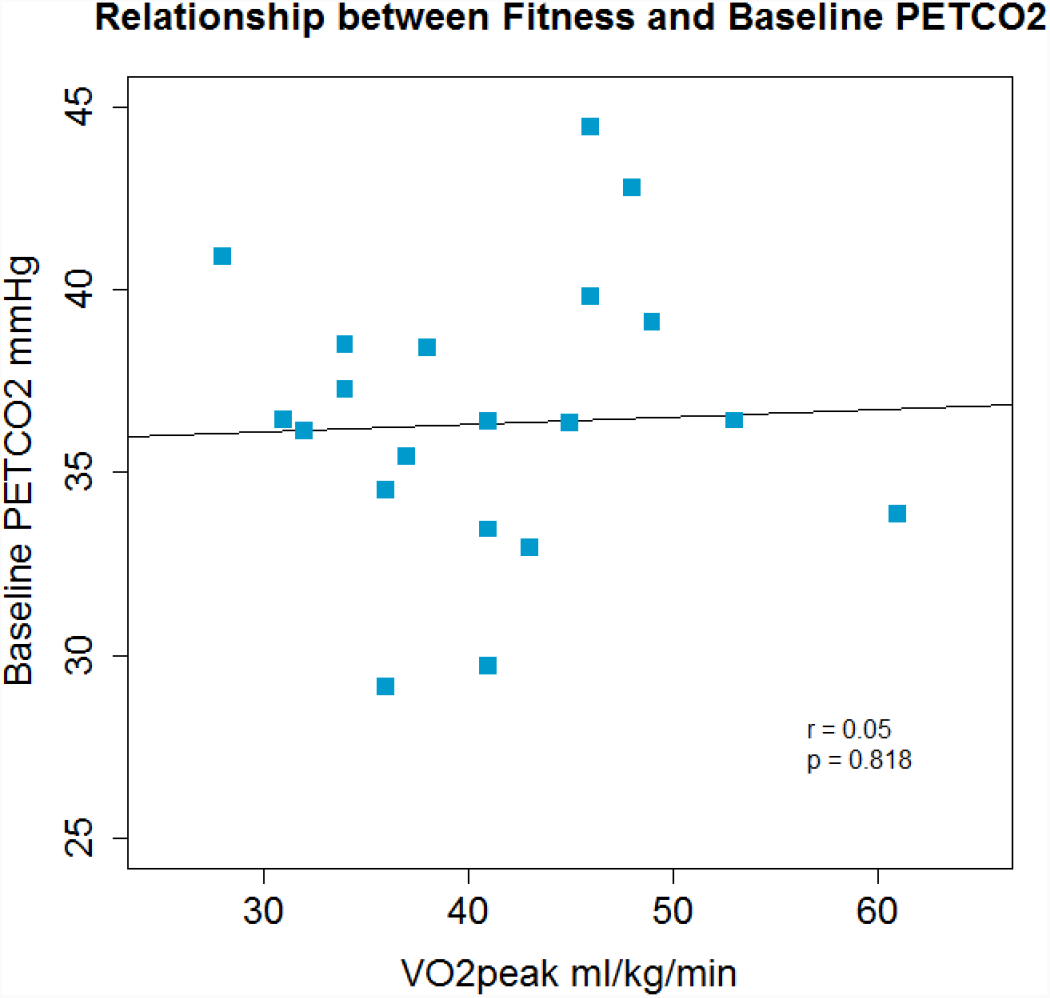
Aerobic fitness was not a predictor of baseline P_ET_CO_2_.

**Figure S3.**
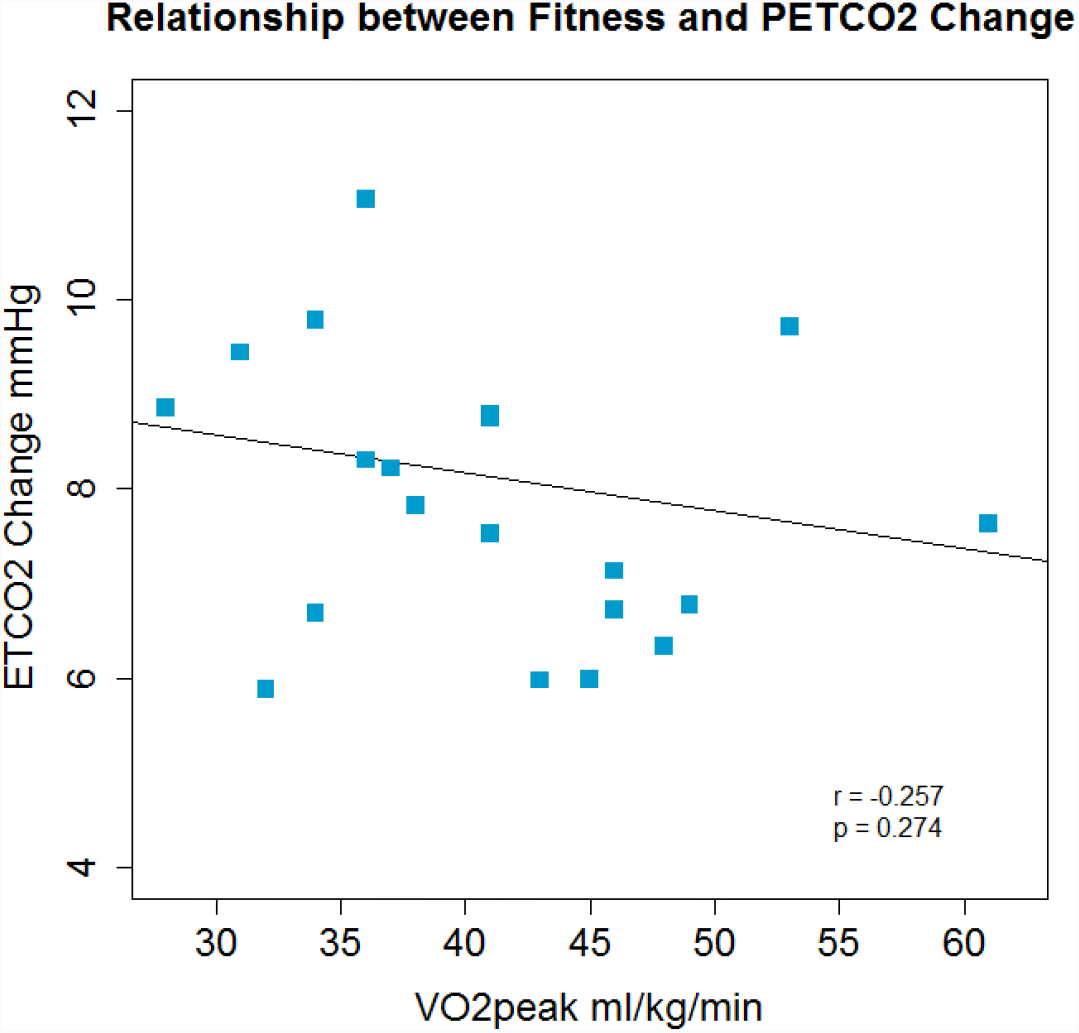
Aerobic fitness was not a predictor of ΔP_ET_CO_2_ during hypercapnia.

**Figure S4.**
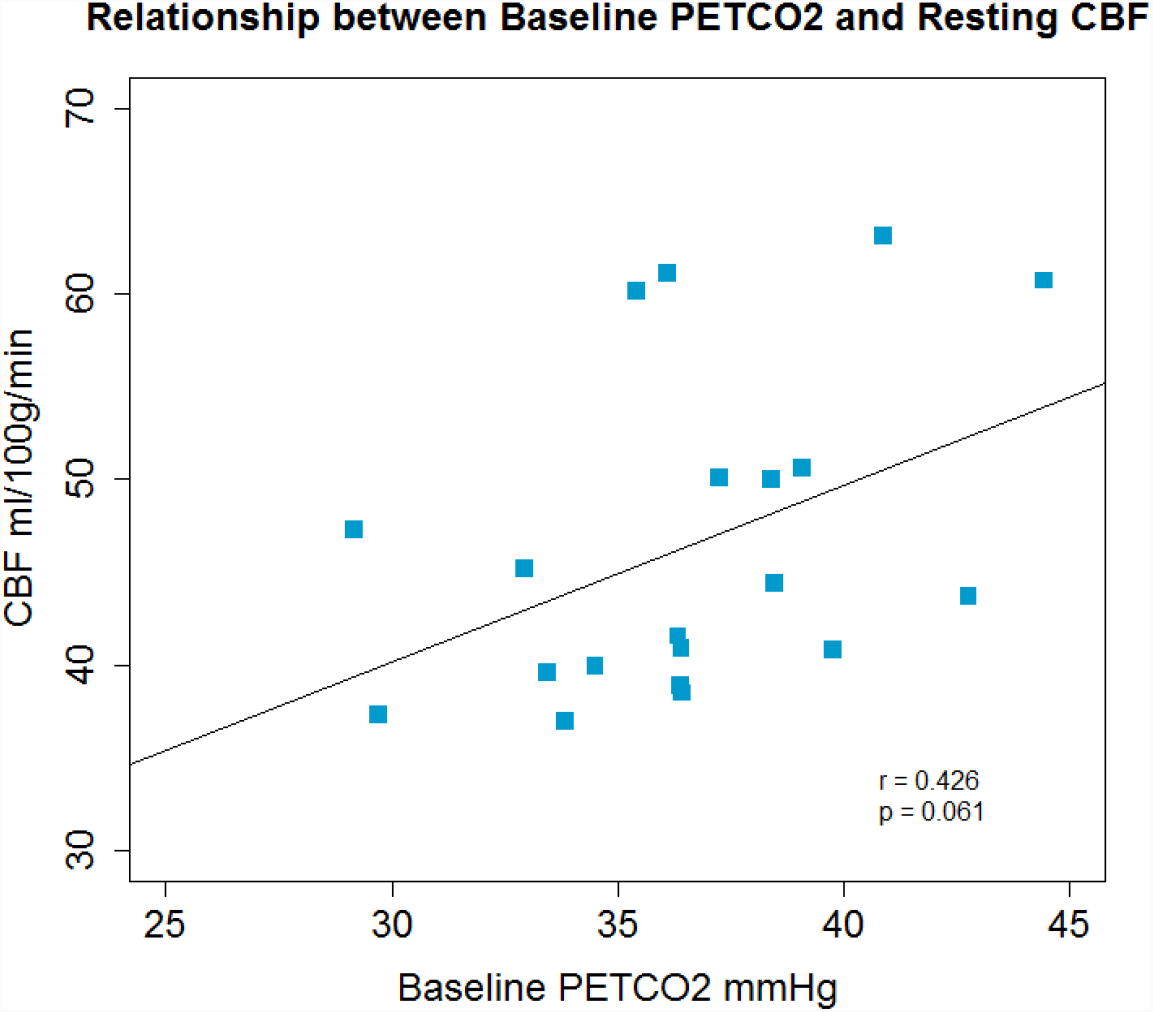
There was a non-significant positive association between resting CBF and baseline P_ET_CO_2_.

**Table S1.** Group average and standard deviation for each cognitive test.

| Cognitive Test | Mean (sd) |
| --- | --- |
| Conners Continuous Performance (d') | 43.6 (7.6) |
| SDMT | 62 (10) |
| Trail Making B | 51.3 (28.5) (s) |
| SCOLP (max 100) | 81 (17) |
| Digit Span (LDSF) | 6.8 (1.4) |
| Letter Fluency | 45.1 (8.6) |
| HADS A | 3.75 (2.2) |
| HADS D | 0.7 (0.9) |

**Table S2.**
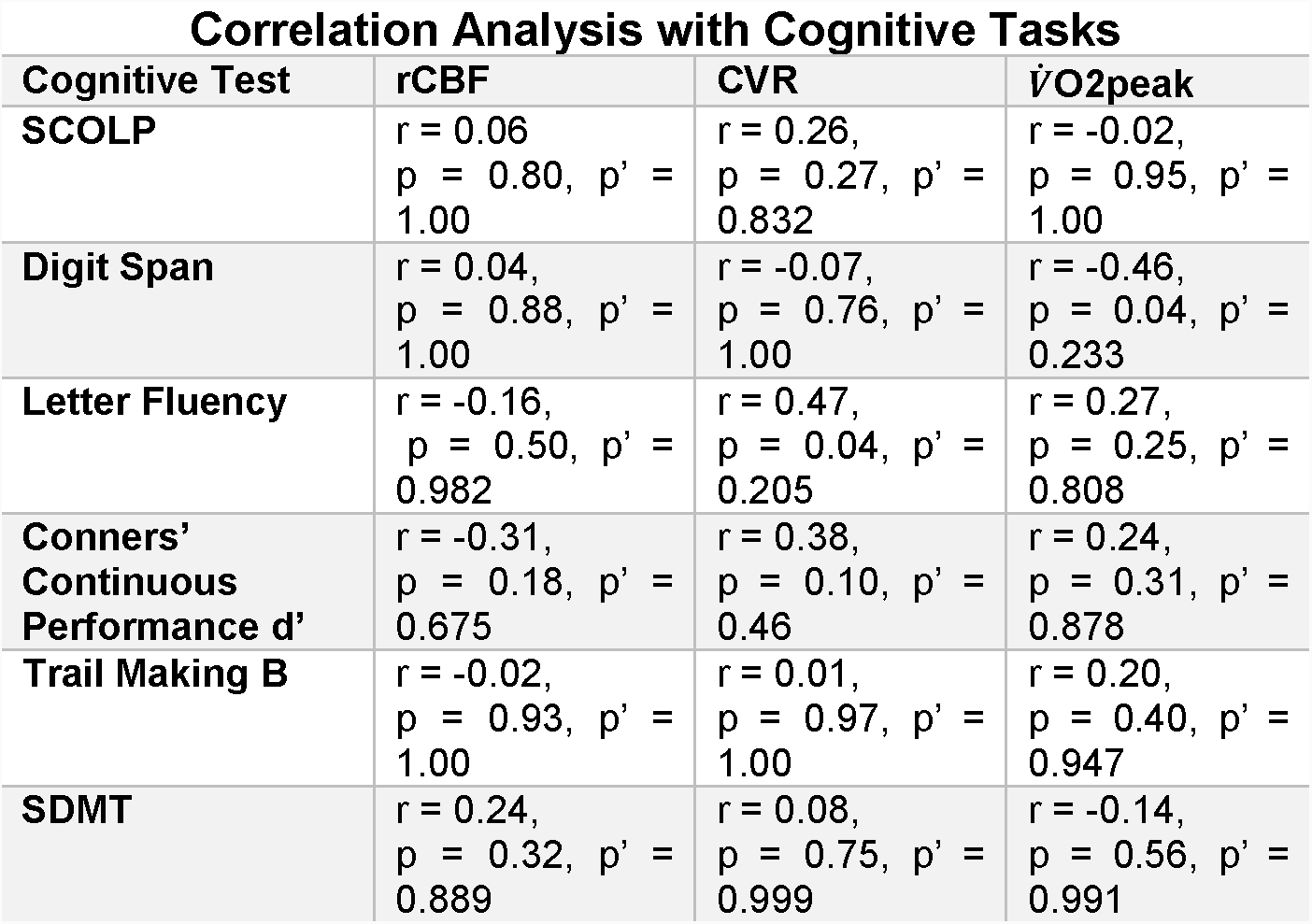
Correlations between cognitive task performance and fitness as well as CBF and CVR. RCBF = Resting CBF. p′ = corrected p value.

## Notes

This work was supported by the Wellcome Trust [grant numbers [506285] and [200804/Z/16/Z] and The Waterloo Foundation.

